# Genome-wide Methylation Patterns Under Caloric Restriction in *Daphnia magna*

**DOI:** 10.1101/278408

**Authors:** Jack Hearn, Marianne Pearson, Mark Blaxter, Philip Wilson, Tom J. Little

**Author notes:** Email addresses: JH MP MB PW TJL.

## Abstract

The degradation of epigenetic control with age is associated with progressive diseases of ageing, including cancers, immunodeficiency and diabetes. Reduced caloric intake slows the effects of aging and age-related diseases, a process likely to be mediated by the impact of caloric restriction on epigenetic factors such as DNA methylation. We used whole genome bisulphite sequencing to study how DNA methylation patterns change with diet in a small invertebrate, the crustacean *Daphnia magna*. *Daphnia* show the classic response of longer life under CR, and they reproduce clonally, which permits the study of epigenetic changes in the absence of genetic variation. Global CpG methylation was 0.7-0.9%, and there was no difference in overall methylation levels between normal and calorie restricted replicates. However, 453 regions were differentially methylated (DMRs) between the normally fed and calorie restricted (CR) replicates. Of these 61% were hypomethylated in the CR group, and 39% were hypermethylated in the CR group. Gene Ontogeny (GO) term enrichment of hyper and hypo-methylated genes showed significant over- and under-representation in three molecular function terms and four biological process GO terms. Notable among these were kinase and phosphorylation activity, which have a well-known functional link to cancers.

## Introduction

Epigenetic modifications play a key role in maintaining gene expression and organismal development. This is particularly evident when epigenetic control degrades, resulting in progressive diseases in humans, including cancers, immunodeficiency and diabetes [1]. The degradation of epigenetic control with age is proposed to occur in a drift-like process. One mechanism that may rescue age-related epigenetic dysregulation is caloric restriction (CR): reduced caloric intake without malnutrition or loss of nutrients. CR slows the effects of aging and postpones the development of age-related diseases [2–5]. In rhesus monkeys and mice, CR of 30% and 40% respectively appears to reduce epigenetic drift in methylation and increases lifespan, which in rodents can be an extension of up to 50% [6]. Similar results have been seen in yeast, spiders, worms, fish and non-human primates [4,7,8]. CR may also delay a spectrum of diseases such as cancer, kidney disease, autoimmune disease and diabetes [9–11], as well as neurodegenerative diseases [12,13].

DNA Methylation, a reversible covalent modification that regulates gene expression, is the best-studied epigenetic mechanism. DNA Methylation of cytosines occurs when DNA methyltransferase enzymes (DNMTs) transfer a methyl group onto cytosine [14] to create a 5-methylcytosine. Most commonly at a cytosine immediately followed by guanine (CpG site). There are three DNMT enzymes: DNMT3 establishes methylation *de novo*, DNMT1 maintains methylation, and DNMT2, which has no known role in DNA methylation. A reduction in expression levels of DNMT enzymes is associated with ageing, leading to a global loss of genomic methylation [15]. In mammals around 70% of CpGs are methylated [16], however in invertebrates the rate in species sampled to date is lower, from 0% in flies to 15% in the oyster *Crassostrea gigas* [17,18]. The model crustaceans *Daphnia magna* and *Daphnia pulex* (Arthropoda: Crustacea) have genomic CpG methylation of 0.52% and 0.7% respectively [19].

CpG methylation can increase or decrease gene expression dependent on the location of the methylation. In promoter regions, which can be rich in CpGs and are known as CpG islands, it represses expression of the gene. Further to this, many CpG islands are also enriched for permissive chromatin modification, which condenses the structure of chromatin and further prevents transcription. In contrast, methylation of gene bodies leads to an increase in expression of the effected gene. Invertebrates have few CpG Islands, and methylation predominantly occurs in gene bodies, and is enriched in exonic sequence [20–25], where it may enhance transcription or mediate alternative splicing [25–27]. Interestingly, in silkworms there is no correlation between methylation in promoters and gene expression [28], suggesting invertebrates and vertebrates differ in their usage of CpG methylation.

The relationship between diet and CpG methylation, and subsequent impact on ageing and health, is well established. Indeed, DNA methylation may be a predictor of biological age. CR in mice caused a two-year difference in biological (0.8) versus chronological (2.8) age, while in rhesus monkeys CR resulted in a biological age of 20 years for monkeys of chronologically aged 27 years [29]. Specific examples of a diet by methylation interaction include the expression of DNMTs which have elevated expression in response to CR in cancer cells, which counteracts the global hypomethylation [30] observed during ageing. CR also causes a reduction in lipid metabolism gene expression by DNA methylation of gene bodies in mouse livers [31]. As a result, older mice undergoing CR were protected from fatty degeneration, visceral fat accumulation, and hepatic insulin resistance compared to controls. In rats and monkeys short-term CR in older individuals ameliorates the effects of ageing with respect to disease markers, oxidative stress and damage, and increases the expression of longevity related genes [32,33]. The reverse is seen in obesity-like phenotypes in rodent models. For example, in Agouti mice, the agouti viable yellow metastable epiallele (A^vy^) interacts with an upstream retrotransposon intracisternal A particle (IAP) [34]. Unmethylated IAP results in yellow mice and negative health effects associated with obesity, whereas methylation at IAP results in brown healthy mice [35]. Waterland et al (2003)[36] demonstrated the that supplementing mothers with folic acid, vitamin B12, choline and betaine shifted the offspring of obesity phenotype mice to smaller, brown mice indicative of increased methylation at IAP.

Our work aims to determine if an experimentally controlled nutritional environment directs changes in methylation status in a small invertebrate, the crustacean *Daphnia magna*. We do this by whole genome bisulphite sequencing of CR and normally-fed (NF) replicates, identifying and characterizing regions of differential methylation. *Daphnia* show the classic response of longer life under CR and strong maternal effects; the offspring of calorie-restricted mothers being larger and more resistant to pathogens than their counterparts from better fed mothers. Provisioning of offspring, e.g. with carbohydrates, protein or fats, is one explanation for these maternal-effect phenotypes, and epigenetic processes, such as methylation, are also potentially key regulators in these plastic responses to fluctuating environments.

*Daphnia* have many attributes that make them favourable for epigenetic study. First, they reproduce clonally, which permits the study of epigenetic changes in the absence of genetic variation. This also allows powerful study of genetic variation: clonal replicates are equivalent to identical twin studies, but with an experimentally chosen number of uplets. CpG-based methylation occurs in *Daphnia*, its genome encodes all three DNMT enzymes orthologous to mammalian DNMT enzymes, and the global methylation pattern changes in response to environmental factors [19,37–39].

## Results

### Methylated sites prediction

Trimmed bisulphite-converted reads aligned to the genome using Bismark [40] exhibited lower mapping efficiencies than standard short-read alignments typical of WGBS [41], with 20-32% or reads not aligning to the reference genome (read filtering and coverages per replicate Table 1, Bismark report outputs Supplementary File 1). Of the aligned reads, 29-38% of reads were discarded as PCR duplicates, and 6-10% of the remainder contained predicted CHH or CHG methylated sites which were also removed from analyses. This resulted in replicate average read coverages of 8-12-fold (read filtering and coverages per replicate Table 1, Bismark report outputs Supplementary File 1). Global CpG methylation was 0.7-0.9% in all samples, and no difference in methylation levels is observed between normal and calorie restricted replicates. Removal of polymorphic CpG sites, where a polymorphic C/T can be miscalled as methylated C, using variants predicted from the bisulphite-unconverted data had little effect on the total number of sites (table 2). After filtering 99.2% of sites were retained per replicate on average per replicate, with average total sites across replicates going from 6.9 million to 6.85 million. Hierarchical clustering of replicates by methylation status in methylKit [42] demonstrated that mother has a stronger effect on global methylation status than nutritional treatment (Figure 1).

**Table 1:** Read sequenced per replicate for both converted and unconverted libraries. Alignments analysed is number of read-pairs per bisulphite converted library aligned by Bismark; deduplicated is number of read pairs after removal of PCR duplicates; non CpG filtered is read-pairs remaining after removal of CHH and CHG containing reads; bases remaining are number of bases left for CpG methylation prediction in Bismark; average coverage is for filtered bases at a D. magna genome size of 240 megabases; CpG % methylated is the genome wide percentage of CpG methylation. Diet, H: normal food, L: caloric restriction, code: combination of diet and mother.

| Diet | Mother | Code | Converted pairs | Unconverted pairs | Alignments analysed | Deduplicated mapped reads | non CpG filtered reads | Bases remaining | Average coverage | CpG % methylated |
| --- | --- | --- | --- | --- | --- | --- | --- | --- | --- | --- |
| H | 5 | 5H | 25601600 | 14305707 | 18634549 | 12764776 | 12026017 | 2320675218 | 9.67 | 0.7 |
| H | 7 | 7H | 29785044 | 132653916 | 21608960 | 14878231 | 13894067 | 2693793276 | 11.22 | 0.8 |
| H | 8 | 8H | 26405910 | 20875768 | 19223306 | 12837344 | 12060578 | 2312369294 | 9.63 | 0.7 |
| H | 9 | 9H | 29296515 | 12172764 | 21000058 | 14155807 | 13242208 | 2622392293 | 10.93 | 0.8 |
| H | 16 | 16H | 34120276 | 15802459 | 23704762 | 15726061 | 14675418 | 2914262560 | 12.14 | 0.8 |
| H | 22 | 22H | 34878128 | 23164024 | 15908472 | 15908472 | 14830368 | 2919581788 | 12.16 | 0.8 |
| L | 5 | 5L | 22562863 | 21579618 | 14154784 | 10164439 | 9447332 | 1871992335 | 7.80 | 0.7 |
| L | 7 | 7L | 21248753 | 22953919 | 15063166 | 10412350 | 9743258 | 1937024597 | 8.07 | 0.7 |
| L | 8 | 8L | 31610110 | 24339207 | 21765927 | 14712500 | 13701673 | 2657817364 | 11.07 | 0.8 |
| L | 9 | 9L | 34684920 | 20342139 | 23809968 | 15706084 | 14477225 | 2870659128 | 11.96 | 0.8 |
| L | 16 | 16L | 29775932 | 19257227 | 19183633 | 13586009 | 12679198 | 2523165965 | 10.51 | 0.8 |
| L | 22 | 22L | 32239102 | 32281921 | 14083083 | 14083083 | 12718680 | 2438782524 | 10.16 | 0.9 |

**Table 2:** CpG sites remaining in each replicate after filtering of polymorphic sites demonstrating minimal effect of this filtering. Polymorphic sites were identified from unconverted libraries against a reference genome converted to Clone 32. Diet, H: normal food, L: caloric restriction, code: sample code.

| Diet | Mother | Code | Sites before SNP filtering | After SNP filtering | % Remaining |
| --- | --- | --- | --- | --- | --- |
| H | 5 | 5H | 6298891 | 6247063 | 99.18 |
| H | 7 | 7H | 6983214 | 6926563 | 99.19 |
| H | 8 | 8H | 6283291 | 6231498 | 99.18 |
| H | 9 | 9H | 7299503 | 7240944 | 99.20 |
| H | 16 | 16H | 7325180 | 7266539 | 99.20 |
| H | 22 | 22H | 7450910 | 7391057 | 99.20 |
| L | 5 | 5L | 6577176 | 6523622 | 99.19 |
| L | 7 | 7L | 6456008 | 6403288 | 99.18 |
| L | 8 | 8L | 6934926 | 6878855 | 99.19 |
| L | 9 | 9L | 7476524 | 7416772 | 99.20 |
| L | 16 | 16L | 7335517 | 7276624 | 99.20 |
| L | 22 | 22L | 6392175 | 6339761 | 99.18 |

**Figure 1:**
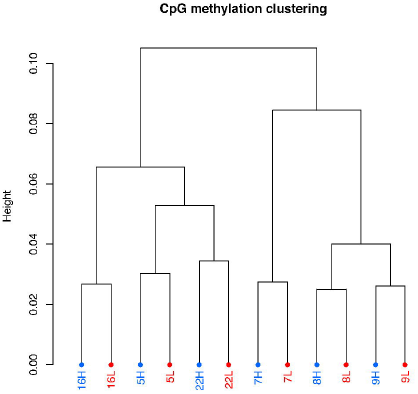
Samples cluster by mother and not by treatment in global CpG similarity. Dendrogram created by ward.D method in methylKit. Number refers to mother from which replicate was derived; H for normal food diet, L for caloric restriction.

### Differential methylation in bsseq

Bsseq [43] testing of differential methylation revealed 453 differentially methylated regions (DMRs) using a t-statistic cutoff of −4.6, 4.6. Of these 278 (61%) were hypomethylated in the CR group versus normal food, and 175 were hypermethylated (39%) in the CR group versus normal food. DMRs were 164 base-pairs (bp) and 127 bp long for CR hypo-and CR hyper-methylated regions and ranged from 10-602 bp in length. There are from three to 20 CpGs per cluster with an average of six. CR hypomethylated DMRs overlapped 357 gene predictions in the *Daphnia magna* geneset, while hypermethylated DMRs overlapped 244. The majority of these also overlapped exonic sequences: 99% (353/357) for CR hypomethylated DMRs and 80% (194/244) for CR hypermethylated DMRs. Only 63 DMRs (14%) do not overlap a predicted gene body at all. The increase in number of DMR containing genes versus total DMRs reflects overlapping/redundant predictions in the *D. magna* genome annotation version 2.4.

### GO term enrichment in methylated genes and DMRs

GO term enrichment was explored using the “weight01” and “classic” algorithms in topGO [44] and molecular function (MF) and biological process (BP) terms are reported. The enrichment analysis (DMRs) using the weight01 algorithm showed significant over and under representation in three molecular function (MF) GO terms and four biological process (BP) terms. The more permissive classic algorithm revealed significant enrichment in twenty-five MF GO terms and twenty BP. These results have been condensed into their most-specific terms and direction of methylation in Table 3 (expanded version, Supplementary File 2). Top-scoring DIAMOND [45] alignment by bit-score against the uniref90 [46] and non-redundant protein databases for each gene associated with an enriched GO terms is reported in Supplementary File 3.

**Table 3:** Functional enrichment of DMRs using the Biological Process and Molecular Function Gene Ontology (GO). Table lists all significant terms identified with the ‘weight01’ algorithm, which accounts for GO term hierarchy, plus any that were only identified with the ‘classic’ algorithm, which ignores hierarchy.

| Biological process |  |  |  |  |
| --- | --- | --- | --- | --- |
| GO ID | GO term | Ancestors | Algorithm | Direction |
| GO:0006468 | protein phosphorylation | 7 | weight01 | **Hyper** |
| GO:0043039 | tRNA aminoacylation | 2 | weight01 | **Hyper** |
| GO:0031167 | rRNA methylation | 4 | weight01 | **Hyper** |
| GO:0022904 | respiratory electron transport chain | 0 | classic | **Hyper** |
| GO:0006644 | phospholipid metabolic process | 1 | classic | Hypo |
| GO:0046486 | glycerolipid metabolic process | 1 | classic | Hypo |
| Molecular function |  |  |  |  |
| GO:0004713 | protein tyrosine kinase activity | 5 | weight01 | **Hyper** |
| GO:0005524 | ATP binding | 12 | weight01 | **Hyper** |
| GO:0042626 | ATPase activity, coupled to transmembran... | 1 | weight01 | Hypo |
| GO:0015405 | P-P-bond-hydrolysis-driven transmembrane... | 1 | classic | Hypo |

## Discussion

### Global methylation and differentially methylated regions

The global CpG methylome of ~0.7%, consistent across replicates, is possibly higher than for the previously sequenced *D. magna* CpG methylome of 0.5% [19], though this earlier study was performed on a different strain and used different data filtering methods [19]. In line with previously observed methylation patterns in arthropods [20,23,24,47], the majority (86%) of DMRs are found in gene bodies. Furthermore, most are present in exonic regions, although more so for hypo-(99%) than hypermethylated (80%) regions. This suggests DNA methylation is regulating expression of targeted genes in *Daphnia* as for other invertebrates [20,47]. Although this requires confirmation by gene expression data, the expectation is that hypermethylated genes are upregulated in expression and hypomethylated are downregulated.

In what follows, we discuss the genes that are associated with the functional enrichment of differentially methylated regions (GO terms, Table 3), giving particular attention to genes whose expression is known to respond to CR, or are linked to progressive disease of ageing or cancers.

### Hypermethylation under CR

Perhaps the most prominent difference between CR and control *Daphnia* is the hypermethylation of protein phosphorylation and tyrosine kinase activity (Table 3, BP, GO:0006468; MF, GO:0004713). These GO terms are associated with genes that include four calcium/calmodulin-dependent protein kinase kinases (CAM-KK). CAM-KKs are involved in regulating cell apoptosis and promote cell survival by activating protein kinase B (Akt) [48], which also has roles in cell-cycle progression. In humans, aberrant expression of CAM-KK is a known factor in several cancers, and is considered a therapeutic target for prostate and stomach cancers [49,50]. CAM-KK responses to CR are less well understood, but can protect against atherosclerosis by activation of AMP-activated protein kinase and sirtuin-1 [51]. Sirtuin proteins are involved in the response to CR and general nutrient sensing [52,53]. CAM-KK upregulation due to hypermethylation under CR thus might activate sirtuin-based responses and subsequent whole organism phenotypes. Further investigation of this might included characterising differences in downstream expression of AMPK, sirtuins, mTORC1 [54] and sirtuin triggered pathways.

The next major hypermethylated GO term, the molecular function term for ATP binding (GO:0005524), is associated with a gene group that includes kinases, ligases, and eleven uniquely occurring genes (Supplementary File 3). Three of these (ABCD I, GMP synthase and The RNA helicase DDX39A), have established links to diet or ageing. ATP-binding cassette sub-family D member 1 (ABCD1), is involved in the catabolism of long chain fatty acids [55]. This suggests upregulation of energy production in our CR lines (though we also find evidence of down-regulation of lipid metabolism, discussed below). Mutations in this gene in humans causes adrenomyeloneuropathy, characterised by an accumulation of unbranched saturated fatty acids [56]. In ABCD1 knockout mice cholesterol levels are higher than in wildtypes, and are unaffected by cholesterol feeding [57]. Most pertinent to CR, however, is the defective antioxidant response correlated with ABCD1 dysfunction [58], because reducing oxidative stress is a proposed mechanism by which CR increases longevity [6,59,60]. GMP synthase expression gradually decreases with age resulting in lower cognitive performance [61]. The RNA helicase DDX39A has no connection to CR, but its overexpression is associated with poor cancer prognosis [62–66].

Also among the eleven uniquely occurring genes associated with the GO term for ATP binding are several genes associated with DNA stability (Supplementary File 3). SMC2, for example, is a component of the condensing complex which organises and condenses chromosomes during mitosis and meiosis [67,68]. SMC2 also acts to repair double-stranded breaks in DNA and maintains ribosomal DNA stability in yeast, dysregulation of which is linked to cancer in humans [67,69]. Although no link has been established between CR and SMC2, CR enhances genomic stability through several pathways including double-strand repair [70]. The condensin complex is a possible mechanism by which this enhancement occurs. The presence of DNA polymerase θ, and the MCM2 component of the MCM2-7 complex, which is essential in initiating DNA replication by unwinding double-stranded DNA, is further evidence of a genomic stability maintenance response of CR. A reduction in expression of the MCM2-7 complex leads to aneuploidy and in mice reduced life-spans due to cancer [71,72]. DNA polymerase θ also acts to repair double-stranded breaks in DNA, and higher expression of this gene is associated with better cancer treatment outcomes [73].

The tRNA aminoacylation ligase (GO:0043039) group (Table 3) is associated with four tRNA ligases (for glutamate, proline, histidine, and phenylalanine). There is no research directly linking these ligase genes to CR, but increased expression via methylation may be a response to low abundance of these amino acids, which indicates that future studies of diet should vary protein availability. Indeed, it may be that protein restriction is more important than overall calorie restriction for longevity (cites). Additionally, fragments of tRNAs called 5’ tRNA halves are a class of signalling molecules that are modulated by CR and ageing in mice [74,75], in which CR ‘rescues’ older 5’ tRNA halves in line with other CR phenotypes. An as yet undiscovered mechanism of diet and ageing could involve tRNA ligases regulating levels of 5’ tRNA halves in response to CR.

Respiratory electron transport chain (GO:0022904; Table 3) contains two proteins: cytochrome b-c1 complex subunit and NADH (Nicotinamide adenine dinucleotide reduced form) dehydrogenase 1 alpha subcomplex subunit. Their methylation may relate to more efficient respiration because of CR to extract maximum energy from food. Interestingly, in yeast, CR is associated with increased longevity due to a reduction in NADH levels because of NADH dehydrogenase activity [76] to create NAD+ (oxidised form). NADH is a competitive inhibitor of yeast sirtuin, leading to its activation on decreased NADH levels [76,77]. This could also be occurring in *Daphnia* under CR if methylation of the NADH gene results in the expected increase in gene expression. The RNA methylation group for hypermethylated genes (GO:0031167) contains two methyltransferase protein 20s (not DNA methyltransferases) which do not have a clear link to CR in the literature.

### Hypomethylation under CR

Phospholipid/glycerolipid metabolism is reduced under CR (Table 3, GO:0006644 and GO:0046486), and both processes are associated with the same genes. These genes are GPI inositol-deacylase, cardiolipin synthase, phosphatidylinositol-glycan biosynthesis class W protein, and phosphatidylserine synthase. Assuming that decreased methylation lowers gene expression, this result is in keeping with previous work on effects of CR on phospholipids. In mice myocardium, phospholipids undergo a reduction in mass and are remodelled when facing CR [78], which is speculated to maximise energy efficiency. The same drop in phospholipids was observed in humans undergoing acute CR [79], and more generally reduces the rick of atherosclerosis and heart disease [80,81]. This is potentially a further common mechanism of response to CR in which DNA methylation is an important component. The molecular function GO term for ATPase activity (GO:0042626) and overlapping P-P-bond-hydrolysis-driven transporters (GO:0015405) include plasma membrane calcium-transporting ATPase, downregulation of which would increase calcium/calmodulin-dependent protein kinase activity within cells by maintaining calcium levels. The remaining genes encode ATP binding protein sub-family B proteins, which pump various substrates out of cells and have no clear links to known CR phenotypes. This may reflect the broad-range of substrates ATP binding protein are able to efflux.

## Conclusion

We have shown that caloric restriction effects the methylation status of a subset of genes, despite the low overall CpG methylation found in *Daphnia*. There is a strong concordance between these results and CR experiments in humans, mice and yeast among other species. We show that hypermethylated genes and processes are in line with upregulation in previous CR and hypomethylated genes with downregulation. Although we have focused on the effect of caloric restriction on DNA methylation status, there are alternative potential epigenetic responses to CR, including small RNAs (sRNAs) and histone modifications. Previously, we established that CR induces differential miRNA expression in *D. magna* under an equivalent experimental design [82], but other sRNAs, for example piRNAs and tsRNAs, could also have a role in CR-dependent gene regulation [83–85]. Histone modifications in response to CR or protein restriction (PR) are known from work on humans, rats and mice [86–89] and are proposed to increase longevity [87] by delaying and repressing ageing-related processes and diseases. Future studies could vary a range of dietary components (overall calories, proteins or fatty acids) and examine the joint effects of a range of epigenetic mechanisms.

## Methods

### Daphnia preparation and experiment

Six replicates of control (i.e. well-fed) *Daphnia magna* were compared to six replicates of caloric restricted *Daphnia* to identify differentially methylated regions. We used a single clone (known to us as Clone 32) of *D. magna*, collected from the Kaimes pond near *Leitholm* in the Scottish Borders [90]. Maternal lines were first acclimatized for three generations. For this, individuals were kept in artificial pond medium at 20°C and on a 12h:12h light:dark cycle and fed 2.5 ×10^6^ cells of the single-celled green algae *Chlorella vulgaris* daily. Following three generations of acclimatisation (detailed in [82]), 40 offspring from each mother were isolated and split to form a replicate. Twenty were fed a normal diet of 5×10^6^ algal cells/day and the remaining twenty that were fed a calorie restricted diet of 1×10^6^ algal cells/day. Each replicate was split and reared in four sub-replicate jars of five *Daphnia* which were subsequently pooled at DNA extraction. Hence, normal food and calorie restricted replicates and were paired by mother and each consisted of twenty *Daphnia* total. The experiment was ended after the birth of 2^nd^ clutch (approximately day 12 of the treatment generation). *Daphnia* were ground by motorized pestle in Digsol and proteinase K and incubated overnight at 37°C and stored at −70°C until DNA extraction.

### DNA extraction and sequencing

DNA was extracted from pooled *Daphnia* per replicate by phenol-chloroform followed by a Riboshredder RNA digestion step and repeat of the phenol-chloroform step. DNA was eluted into 100 ul of TE buffer and quantified by Qubit fluorimeter Sample purity was checked by 260:280 ratio on nanodrop, and DNA integrity was examined by running approximately 35 ng DNA on a 0.8% agarose gel stained with ethidium bromide. Each DNA extraction was split in two for creation of a bisulphite converted library and corresponding bisulphite unconverted library (all steps the same except bisulphite conversion). Thus, 24 libraries were created: 12 bisulphite-converted and 12 corresponding unconverted samples. This was done to identify per replicate mismatches from the reference and remove false positive methylation calls. All libraries were created by Edinburgh Genomics using the Zymogen EZ DNA Methylation-Lightning Kit and Methylseq Library prep Illumina TruSeq DNA Methylation Kit and 125 base pair paired-end sequenced on Illumina HiSeq. Raw read data has been deposited in the European Nucleotide Archive under accession PRJEB24784, (file names and conversion status, Supplementary File 4).

### Quality Assessment and Mapping

Before aligning reads to the *D. magna* reference genome (version 2.4 downloaded from: http://arthropods.eugenes.org/EvidentialGene/daphnia/daphnia_magna/Genes/earlyaccess/), the reference was converted to clone 32 as for Hearn et al (2018) [82]. This was to increase mapping efficiency, and accuracy of the analysis, by reducing polymorphism between the reference (assembled from a different clone) and our data.

Reads from both bisulphite converted and unconverted libraries were trimmed of the first and last nine bases of every read using TrimGalore! (version 0.4.1) [91] after initial inspection of Bismark m-bias plots. TrimGalore! was also used to remove base calls with a Phred score of 20 or lower, adapter sequences, and sequences shorter than 20 bp. FastQC 0.11.4 [92] was used to before and after quality control to inspect the data. Bisulphite calls were made with Bismark 0.16.3 [40]. Bismark alignments were performed with options “– score_min L,0,-0.6”. PCR duplicates were removed using deduplicate_bismark script. Bismark reports indicated that libraries were not fully bisulphite converted and raw methylation calls were approximately 3% for CpG, CHH and CHG sites. Previous research has shown that CHH and CHG methylation is negligible in the *D. magna* genome [19]. As a result, we used the filter_non_conversion script to remove reads that contain either CHH and CHG methylation sites as diagnostic of a non-bisulphite converted read. Finally, methylated sites were identified using bismark_methylation_extractor and reports created with bismark2report.

Further variants were predicted per replicate using the unconverted library reads by following the GATK pipeline [93,94] and converted strain 32 reference sequence. One round of base quality score recalibration was sufficient using variants previously identified from strain 32. Sites with single nucleotide polymorphisms at methylated positions were removed from the analysis using BEDtools [95].

### Differential Methylation Analysis

All analyses were performed using methylation calls from the bisulphite converted libraries only. Hierarchical sample clustering of genome-wide methylation patterns across replicates was generated using methylKit [42]. The six-normal food-and six calorie restricted replicates were then compared using bsseq [43] Bioconductor package in R to identify regions of differential methylation. We ran bsseq with a paired t-test to control for the batch effect of mother on methylation. DMRs were selected using t-statistic cutoff of −4.6 and 4.6, a greater than 0.1 average difference in methylation between groups, and at least three methylated CpGs. Genes overlapping DMRs were identified using the *D. magna* version 2.4 genome annotation file and BEDTools. This list of overlapping genes was used as a basis for functional enrichment analysis.

### Functional Enrichment Analysis

Enriched GO (gene ontology) terms were identified using topGO [44] (weight01 and classic algorithm) and molecular function (MF) and biological process (BP) GO terms are reported. We tested if differentially methylated genes were enriched for specific GO terms against a database of GO terms for all the genes in the *Daphnia magna* genome. Differentially methylated regions were also split into hyper and hypo-methylated in calorie restricted categories to test for direction-specific enrichment. *D. magna* GO terms were downloaded from: http://arthropods.eugenes.org/EvidentialGene/daphnia/daphnia_magna/Genes/function/cddrps-dapmaevg14.gotab2. Fisher’s exact test with an *a* of 0.01 was used to identify enriched genes, and no p-value correction was applied as per topGO author recommendation. The group of genes present in each hyper- or hypo- methylated enriched GO terms were extracted and blast [96] searched against uniref90 and the non-redundant protein databases downloaded January 18^th^ 2018 using DIAMOND [45].

## Acknowledgements

We thank Peter Fields, University of Basel and Don Gilbert, Indiana University for providing scripts, advice, and data on *Daphnia magna* genome annotation. This research was funded by Leverhulme Trust grant RPG-2015-406.

---

Supplementary File 1. MultiQC report in html format reporting general statistics from the Bismark alignment process for all replicates. Including alignment rates, deduplication effect, overall cytosine methylation and m-bias plot. This plot shows average methylation level per position across reads, demonstrating minimal bias at 5’ and 3’ reads after trimming of first and last nine base pairs of each read.

Supplementary File 2. Functional enrichment of DMRs using the Biological Process Gene Ontology (GO), Most specific term listed first, and is identified by the weight01 algorithm, indented beneath are GO term description using topGo to test for enrichment blog post from the same hierarchy, identified by the classic algorithm.

Supplementary File 3. DIAMOND aligner results in blast output format six for best match to uniref 90 and non-redundant protein databases of each *D. magna* gene associated with an enriched GO term under the weight01 algorithm. Column one is GO term, column two is *D. magna* gene, columns three-thirteen correspond to standard blast tabular output, and column fourteen is the description of the best-matching hit for that gene.

Supplementary File 4. Raw read files in the European Nucleotide Archive for each replicate, including ENA alias, bisulphite converted or unconverted status, unique identification for that replicate, mother identification and treatment: H is normal food and L is caloric restriction.

## References

[1] Egger G, Liang G, Aparicio A, et al. Epigenetics in human disease and prospects for epigenetic therapy. Nature. 2004;429:457.

[2] Cooper TM, Mockett RJ, Sohal BH, et al. Effect of caloric restriction on life span of the housefly, *Musca domestica*. FASEB J. 2004;18:1591–1593.

[3] Forster MJ, Morris P, Sohal RS. Genotype and age influence the effect of caloric intake on mortality in mice. FASEB J. 2003;17:690–692.

[4] Colman RJ, Anderson RM, Johnson SC, et al. Caloric restriction delays disease onset and mortality in rhesus monkeys. Science (80-.). 2009;325:201–204.

[5] McCay CM, Crowell MF, Maynard LA. The effect of retarded growth upon the length of life span and upon the ultimate body. J. Nutr. 1935;10:63–79.

[6] Sohal RS, Weindruch R. Oxidative Stress, Caloric Restriction, and Aging. Science (80-.). 1996;273:59 LP–63.

[7] Weindruch R. The retardation of aging by caloric restriction: studies in rodents and primates. Toxicol. Pathol. 1996;24:742–745.

[8] Weindruch R, Walford RL. Retardation of aging and disease by dietary restriction. CC Thomas; 1988.

[9] Fernandes G, Yunis EJ, Good RA. Suppression of adenocarcinoma by the immunological consequences of calorie restriction. Nature. 1976;263:504.

[10] Sarkar NH, Fernandes G, Telang NT, et al. Low-calorie diet prevents the development of mammary tumors in C3H mice and reduces circulating prolactin level, murine mammary tumor virus expression, and proliferation of mammary alveolar cells. Proc. Natl. Acad. Sci. 1982;79:7758–7762.

[11] Kubo C, Johnson BC, Good RA. A crucial influence of total calorie intake on autoimmune-prone mice: Influence of diets of grossly different composition on immunologic functions. Fed. Proc. 1984.

[12] Duan W, Mattson MP. Dietary restriction and 2-deoxyglucose administration improve behavioral outcome and reduce degeneration of dopaminergic neurons in models of Parkinson’s disease. J. Neurosci. Res. 1999;57:195–206.

[13] Zhu H, Guo Q, Mattson MP. Dietary restriction protects hippocampal neurons against the death-promoting action of a presenilin-1 mutation. Brain Res. 1999;842:224–229.

[14] Bestor TH. Activation of mammalian DNA methyltransferase by cleavage of a Zn binding regulatory domain. EMBO J. 1992;11:2611–2617.

[15] Pal S, Tyler JK. Epigenetics and aging. Sci. Adv. 2016;2:e1600584.

[16] Ehrlich M, Gama-Sosa MA, Huang L-H, et al. Amount and distribution of 5-methylcytosine in human DNA from different types of tissues or cells. Nucleic Acids Res. 1982;10:2709–2721.

[17] Bewick AJ, Vogel KJ, Moore AJ, et al. Evolution of DNA methylation across insects. Mol. Biol. Evol. 2017;34:654–665.

[18] Olson CE, Roberts SB. Genome-wide profiling of DNA methylation and gene expression in *Crassostrea gigas* male gametes. Front. Physiol. 2014;5 JUN:1–7.

[19] Asselman J, De Coninck DIM, Pfrender ME, et al. Gene Body Methylation Patterns in *Daphnia* Are Associated with Gene Family Size. Genome Biol. Evol. 2016;8:1185–1196.

[20] Zemach A, McDaniel IE, Silva P, et al. Genome-wide evolutionary analysis of eukaryotic DNA methylation. Science (80-.). 2010;328:916–919.

[21] Sarda S, Zeng J, Hunt BG, et al. The evolution of invertebrate gene body methylation. Mol. Biol. Evol. 2012;29:1907–1916.

[22] Feng S, Cokus SJ, Zhang X, et al. Conservation and divergence of methylation patterning in plants and animals. Proc. Natl. Acad. Sci. 2010;107:8689–8694.

[23] Lyko F, Foret S, Kucharski R, et al. The honey bee epigenomes: differential methylation of brain DNA in queens and workers. PLoS Biol. 2010;8:e1000506.

[24] Bonasio R, Li Q, Lian J, et al. Genome-wide and caste-specific DNA methylomes of the ants *Camponotus floridanus* and *Harpegnathos saltator*. Curr. Biol. 2012;22:1755–1764.

[25] Suzuki MM, Kerr ARW, De Sousa D, et al. CpG methylation is targeted to transcription units in an invertebrate genome. Genome Res. 2007;17:625–631.

[26] Flores K, Wolschin F, Corneveaux JJ, et al. Genome-wide association between DNA methylation and alternative splicing in an invertebrate. BMC Genomics. 2012;13:480.

[27] Li-Byarlay H, Li Y, Stroud H, et al. RNA interference knockdown of DNA methyl-transferase 3 affects gene alternative splicing in the honey bee. Proc. Natl. Acad. Sci. 2013;110:12750–12755.

[28] Xiang H, Zhu J, Chen Q, et al. Single base-resolution methylome of the silkworm reveals a sparse epigenomic map. Nat. Biotechnol. 2010;28:516–520.

[29] Maegawa S, Lu Y, Tahara T, et al. Caloric restriction delays age-related methylation drift. Nat. Commun. 2017;8:1–11.

[30] Li Y, Liu L, Tollefsbol TO. Glucose restriction can extend normal cell lifespan and impair precancerous cell growth through epigenetic control of hTERT and p16 expression. FASEB J. 2010;24:1442–1453.

[31] Hahn O, Grönke S, Stubbs TM, et al. Dietary restriction protects from age-associated DNA methylation and induces epigenetic reprogramming of lipid metabolism. Genome Biol. 2017;18:1–18.

[32] Lane MA, Tilmont EM, De Angelis H, et al. Short-term calorie restriction improves disease-related markers in older male rhesus monkeys (*Macaca mulatta*). Mech. Ageing Dev. 2000;112:185–196.

[33] Kim CH, Lee EK, Choi YJ, et al. Short-term calorie restriction ameliorates genomewide, age-related alterations in DNA methylation. Aging Cell. 2016;15:1074–1081.

[34] Waterland RA, Travisano M, Tahiliani KG. Diet-induced hypermethylation at agouti viable yellow is not inherited transgenerationally through the female. FASEB J. 2007;21:3380–3385.

[35] Wolff GL, Kodell RL, Moore SR, et al. Maternal epigenetics and methyl supplements affect agouti gene expression in Avy/a mice. FASEB J. 1998;12:949–957.

[36] Waterland RA, Jirtle RL. Transposable elements: targets for early nutritional effects on epigenetic gene regulation. Mol. Cell. Biol. 2003;23:5293–5300.

[37] Asselman J, De Coninck DIM, Vandegehuchte MB, et al. Global cytosine methylation in *Daphnia magna* depends on genotype, environment, and their interaction. Environ. Toxicol. Chem. 2015;34:1056–1061.

[38] Vandegehuchte MB, De Coninck D, Vandenbrouck T, et al. Gene transcription profiles, global DNA methylation and potential transgenerational epigenetic effects related to Zn exposure history in Daphnia magna. Environ. Pollut. 2010;158:3323–3329.

[39] Vandegehuchte MB, Kyndt T, Vanholme B, et al. Occurrence of DNA methylation in *Daphnia magna* and influence of multigeneration Cd exposure. Environ. Int. 2009;35:700–706.

[40] Krueger F, Andrews SR. Bismark: a flexible aligner and methylation caller for Bisulfite-Seq applications. Bioinformatics. 2011;27:1571–1572.

[41] Porter J, Sun M an, Xie H, et al. Investigating bisulfite short-read mapping failure with hairpin bisulfite sequencing data. BMC Genomics. 2015;16:S2.

[42] Akalin A, Kormaksson M, Li S, et al. methylKit: a comprehensive R package for the analysis of genome-wide DNA methylation profiles. Genome Biol. 2012;13:R87.

[43] Hansen KD, Langmead B, Irizarry RA. BSmooth: from whole genome bisulfite sequencing reads to differentially methylated regions. Genome Biol. 2012;13:R83.

[44] Alexa A, Rahnenfuhrer J. topGO: Enrichment Analysis for Gene Ontology. Available at: https://bioconductor.org/packages/release/bioc/html/topGO.html. 2016.

[45] Buchfink B, Xie C, Huson DH. Fast and sensitive protein alignment using DIAMOND. Nat. Methods. 2015;12:59.

[46] Suzek BE, Wang Y, Huang H, et al. UniRef clusters: a comprehensive and scalable alternative for improving sequence similarity searches. Bioinformatics. 2014;31:926–932.

[47] Hunt BG, Glastad KM, Yi S V, et al. The Function of Intragenic DNA Methylation: Insights from Insect Epigenomes. Integr. Comp. Biol. 2013;53:319–328.

[48] Song G, Ouyang G, Bao S. The activation of Akt/PKB signaling pathway and cell survival. J Cell Mol Med. 2005;9:59–71.

[49] Racioppi L. CaMKK2: A novel target for shaping the androgen-regulated tumor ecosystem. Trends Mol. Med. 2013;19:83–88.

[50] Subbannayya Y, Syed N, Barbhuiya MA, et al. Calcium calmodulin dependent kinase kinase 2 - A novel therapeutic target for gastric adenocarcinoma. Cancer Biol. Ther. 2015;16:336–345.

[51] Wen L, Chen Z, Zhang F, et al. Ca2+/calmodulin-dependent protein kinase kinase β phosphorylation of Sirtuin 1 in endothelium is atheroprotective. 2013;

[52] Guarente L. Calorie restriction and sirtuins revisited. Genes Dev. 2013;27:2072–2085.

[53] Timmers S, Konings E, Bilet L, et al. Calorie restriction-like effects of 30 days of Resveratrol (resVidaTM) supplementation on energy metabolism and metabolic profile in obese humans. Cell Metab. 2011;14:10.1016/j.cmet.2011.10.002.

[54] Igarashi M, Guarente L. mTORC1 and SIRT1 Cooperate to Foster Expansion of Gut Adult Stem Cells during Calorie Restriction. Cell. 2016;166:436–450.

[55] Baker A, Carrier DJ, Schaedler T, et al. Peroxisomal ABC transporters: functions and mechanism. Biochem. Soc. Trans. 2015;43:959–965.

[56] Dean M, Rzhetsky A, Allikmets R. The human ATP-binding cassette (ABC) transporter superfamily. Genome Res. 2001;11:1156–1166.

[57] Weinhofer I, Forss-Petter S, Kunze M, et al. X-linked adrenoleukodystrophy mice demonstrate abnormalities in cholesterol metabolism. FEBS Lett. 2005;579:5512–5516.

[58] Morita M, Imanaka T. Peroxisomal ABC transporters: Structure, function and role in disease. Biochim. Biophys. Acta - Mol. Basis Dis. 2012;1822:1387–1396.

[59] Lee C-K, Klopp RG, Weindruch R, et al. Gene Expression Profile of Aging and Its Retardation by Caloric Restriction. Science (80-.). 1999;285:1390 LP–1393.

[60] Ungvari Z, Parrado-Fernandez C, Csiszar A, et al. Mechanisms underlying caloric restriction and lifespan regulation: Implications for vascular aging. Circ. Res. 2008;102:519–529.

[61] Domek-Łopacińska KU, Strosznajder JB. Cyclic GMP and Nitric Oxide Synthase in Aging and Alzheimer’s Disease. Mol. Neurobiol. 2010;41:129–137.

[62] Nakata D, Nakao S, Nakayama K, et al. The RNA helicase DDX39B and its paralog DDX39A regulate androgen receptor splice variant AR-V7 generation. Biochem. Biophys. Res. Commun. 2017;483:271–276.

[63] Sugiura T, Nagano Y, Noguchi Y. DDX39, upregulated in lung squamous cell cancer, displays RNA helicase activities and promotes cancer cell growth. Cancer Biol. Ther. 2007;6:957–964.

[64] Kikuta K, Kubota D, Saito T, et al. Clinical proteomics identified ATP-dependent RNA helicase DDX39 as a novel biomarker to predict poor prognosis of patients with gastrointestinal stromal tumor. J. Proteomics. 2012;75:1089–1098.

[65] Kubota D, Okubo T, Saito T, et al. Validation study on pfetin and ATP-dependent RNA helicase DDX39 as prognostic biomarkers in gastrointestinal stromal tumour. Jpn. J. Clin. Oncol. Oxford University Press; 2012. p. 730–741.

[66] Otake K, Uchida K, Ide S, et al. Identification of DDX39A as a potential biomarker for unfavorable neuroblastoma using a proteomic approach. Pediatr. Blood Cancer. 2016;63:221–227.

[67] Wu N, Yu H. The Smc complexes in DNA damage response. Cell Biosci. 2012;2:5.

[68] Strunnikov A V., Jessberger R. Structural maintenance of chromosomes (SMC) proteins: Conserved molecular properties for multiple biological functions. Eur. J. Biochem. 1999;263:6–13.

[69] Je EM, Yoo NJ, Lee SH. Mutational and expressional analysis of SMC2 gene in gastric and colorectal cancers with microsatellite instability. APMIS. 2014;122:499–504.

[70] Heydari AR, Unnikrishnan A, Lucente LV, et al. Caloric restriction and genomic stability. Nucleic Acids Res. 2007;35:7485–7496.

[71] Passerini V, Ozeri-Galai E, MS De Pagter, et al. The presence of extra chromosomes leads to genomic instability. Nat. Commun. 2016;7.

[72] Pruitt SC, Bailey KJ, Freeland A. Reduced Mcm2 Expression Results in Severe Stem/Progenitor Cell Deficiency and Cancer. Stem Cells. 2007;25:3121–3132.

[73] Wood RD, Doublié S. DNA polymerase θ (POLQ), double-strand break repair, and cancer. DNA Repair (Amst). 2016;44:22–32.

[74] Dhahbi JM, Spindler SR, Atamna H, et al. 5’ tRNA halves are present as abundant complexes in serum, concentrated in blood cells, and modulated by aging and calorie restriction. BMC Genomics. 2013;14:1–14.

[75] Dhahbi JM. Circulating small noncoding RNAs as biomarkers of aging. Ageing Res. Rev. 2014;17:86–98.

[76] Lin SJ, Ford E, Haigis M, et al. Calorie restriction extends yeast life span by lowering the level of NADH. Genes Dev. 2004;18:12–16.

[77] Imai S, Guarente L. It takes two to tango: NAD+ and sirtuins in aging/longevity control. npj Aging Mech. Dis. 2016;2:16017.

[78] Han X, Cheng H, Mancuso DJ, et al. Caloric restriction results in phospholipid depletion, membrane remodeling, and triacylglycerol accumulation in murine myocardium. Biochemistry. 2004;43:15584–15594.

[79] Collet TH, Sonoyama T, Henning E, et al. A Metabolomic Signature of Acute Caloric Restriction. J. Clin. Endocrinol. Metab. 2017;102:4486–4495.

[80] Fontana L, Meyer TE, Klein S, et al. Long-term calorie restriction is highly effective in reducing the risk for atherosclerosis in humans. Proc. Natl. Acad. Sci. U. S. A. 2004;101:6659–6663.

[81] Heilbronn LK, Ravussin E. Calorie restriction and aging: review of the literature and implications for studies in humans. Am. J. Clin. Nutr. 2003;78:361–369.

[82] Hearn J, Chow FW-N, Barton H, et al. *Daphnia magna* microRNAs respond to nutritional stress and aging but are not transgenerational. Mol. Ecol. 2018;In Press.

[83] Chen Q, Shi J, Peng H, et al. Sperm tsRNAs contribute to intergenerational inheritance of an acquired metabolic disorder. Science (80-.). 2016;351.

[84] Brennecke J, Malone CD, Aravin AA, et al. An epigenetic role for maternally inherited piRNAs in transposon silencing. Science (80-.). 2008;322:1387–1392.

[85] Ashe A, Sapetschnig A, Weick EM, et al. PiRNAs can trigger a multigenerational epigenetic memory in the germline of *C. elegans*. Cell. 2012;150:88–99.

[86] Gong H, Qian H, Ertl R, et al. Histone modifications change with age, dietary restriction and rapamycin treatment in mouse brain. Oncotarget. 2015;6:15882–15890.

[87] Li Y, Daniel M, Tollefsbol TO. Epigenetic regulation of caloric restriction in aging. BMC Med. 2011;9:98.

[88] Sohi G, Marchand K, Revesz A, et al. Maternal Protein Restriction Elevates Cholesterol in Adult Rat Offspring Due to Repressive Changes in Histone Modifications at the Cholesterol 7α-Hydroxylase Promoter. Mol. Endocrinol. 2011;25:785–798.

[89] Sharma R, Nakamura A, Takahashi R, et al. Carbonyl modification in rat liver histones: Decrease with age and increase by dietary restriction. Free Radic. Biol. Med. 2006;40:1179–1184.

[90] Auld SKJR, Hall SR, Housley Ochs J, et al. Predators and patterns of within-host growth can mediate both among-host competition and evolution of transmission potential of parasites. Am. Nat. 2014;184:S77–S90.

[91] Krueger F. Trim galore. A wrapper tool around Cutadapt FastQC to consistently apply Qual. Adapt. trimming to FastQ files. 2015.

[92] Andrews S. FastQC: A quality control application for high throughput sequence data. Available at: http://www.bioinformatics.babraham.ac.uk/projects/fastqc. 2010.

[93] Van Der Auwera GA, Carneiro MO, Hartl C, et al. From FastQ data to high confidence varant calls: the Genome Analysis Toolkit best practices pipeline. Curr Protoc Bioinforma. 2014.

[94] McKenna A, Hanna M, Banks E, et al. The Genome Analysis Toolkit: a MapReduce framework for analyzing next-generation DNA sequencing data. Genome Res. 2010;20:1297–1303.

[95] Quinlan AR, Hall IM. BEDTools: a flexible suite of utilities for comparing genomic features. Bioinformatics. 2010;26:841–842.

[96] Altschul SF, Madden TL, Schäffer AA, et al. Gapped BLAST and PSI-BLAST: a new generation of protein database search programs. Nucleic Acids Res. 1997;25.

